# Chromosome-scale genome assembly of multinucleate heterokaryotic *Rhizoctonia solani* AG1-IB

**DOI:** 10.64898/2026.11.25.769220

**Authors:** Jackson B. Cooper, Mary Anna Carbone, Marc A. Cubeta, James B. White, Rushabh Shah, Ignazio Carbone

## Abstract

*Rhizoctonia solani* AG1-IB is an economically important fungal pathogen of turfgrass and agricultural crops, yet available genome assemblies remain fragmented or represent collapsed or consensus sequences that obscure haplotype-specific variation, limiting investigations of genome evolution and heterokaryosis. Here, we present a chromosome-scale, haplotype-aware genome assembly of AG1-IB isolate S1W6 generated using Oxford Nanopore sequencing, Hi-C scaffolding, and short-read polishing. *De novo* assembly with PECAT produced two haplotype-aware nuclear genome assemblies, with BUSCO completeness greater than 96% and nuclear genome N50 values exceeding 2 Mb. Comparative analyses revealed substantial haplotype divergence, with over 600,000 SNPs and 80,000 indels between haplotypes with 97.75% average nucleotide identity separating assemblies. FastANI analyses showed comparable genetic divergence between Hap1 and Hap2 assemblies of S1W6 to that observed among independently collected AG1-IB isolates from different plant hosts and geographic regions. Despite extensive nucleotide variation, Gene Ontology profiles, predicted effector repertoires, carbohydrate-active enzymes, and biosynthetic gene cluster composition were conserved between assemblies. In contrast, homeodomain (HD) mating-type loci were variable. Hap2 contained four HD genes (*W-HD1*, *W-HD2*, *E-HD1*, and *E-HD2*), whereas Hap1 lacked *E-HD1*, and shared divergent HD amino acid sequences. A *STE3* pheromone receptor gene on chromosome 15 was highly conserved in both haplotypes. This study provides the first chromosome-scale, haplotype-resolved genome resource for *R. solani* AG1-IB and demonstrates that heterokaryotic individuals can harbor genomic variation approaching the levels observed among genetically diverse isolates of AG1-IB. These findings highlight the importance of haplotype-aware assemblies for understanding genome evolution and mating-type diversity in heterokaryotic fungal plant pathogens.

## 1. Introduction

*Rhizoctonia solani* comprises a genetically diverse species complex subdivided into distinct anastomosis groups (AGs) defined by hyphal somatic (in)compatibility (Cubeta and Vilgalys, 1997). The predominantly asexual life history of *R. solani* consists primarily of vegetative mycelium and sclerotia that persist in infested soil and plant debris limiting the generation of genetic diversity through that achieved via sexual recombination and reproduction in some species of *Rhizoctonia*. Despite this constraint, members of the *R. solani* species complex collectively infect more than 250 species of plants across diverse environmental conditions, with substantial host specialization occurring among certain AGs (Ajayi-Oyetunde et al., 2019; Senapati et al., 2022). This contrast raises questions about how genetic diversity is generated and maintained to support adaptation across diverse hosts and environments. The economic importance of *R. solani* is underscored by its occurrence on many major staple food and fiber crops (Anderson et al., 2017). While comparative genomics has revealed substantial genetic divergence among AGs (Anderson et al., 2017; Lee et al., 2021; Li et al., 2021; Wibberg et al., 2021), the mechanism(s) that generate diversity and their relative contributions to population structure within individual AGs remain poorly understood.

Although the reproductive biology of *R. solani* is typically asexual, increasing evidence suggests that it may be a mixture of asexual and sexual reproduction. Field isolates are multinucleate and can exist as heterokaryons or monokaryons, although the frequency and prevalence of these nuclear genome conditions in natural populations have not been the subject of comprehensive investigation 11/25/26 8:57:00 PM. Population-level studies across multiple AGs have identified signatures consistent with gene flow and recombination (Ajayi-Oyetunde et al., 2019; Bernardes-de-Assis et al., 2009; Ciampi et al., 2008; Ferrucho et al., 2013; Muzhinji et al., 2016), and haploid sexual spores (basidiospores) represent a primary source of inoculum in some *Rhizoctonia* pathosystems (Bartz et al., 2010; Shew and Melton, 1995). Recent population genomic analyses of *R. solani* AG1-IA further demonstrated rapid linkage disequilibrium decay and low inbreeding coefficients providing additional indirect evidence that *R. solani* possesses a mixed reproductive system incorporating both clonality and sexual recombination (Gil et al., 2026). In addition, parasexual processes such as heterokaryosis and haploidization may contribute to genetic diversity, although their relative impact in *R. solani* populations is likely constrained by hyphal somatic recognition systems (Adams, 1996; Pontecorvo et al., 1953; Puhalla and Carter, 1976; Schoustra et al., 2007). Recent genomic studies identified homeodomain (HD) genes in *R. solani* homologous to mating-type genes in other fungi in the Phylum Basidiomycota (Coelho et al., 2017; Li et al., 2021; Pardo et al., 1996). For example, four HD genes, *W-HD1*, *W-HD2*, *E-HD1*, and *E-HD2*, have been identified and reported in the Ceratobasidiaceae, with phylogenetic analyses supporting homology of HD and STE3 mating-type proteins to those found in other Basidiomycota fungi, including *Coprinopsis cinerea*, *Cryptococcus neoformans*, *Puccinia striiforms*, and *Ustilago maydis* (Li et al., 2021), suggesting a potential genetic basis for mating compatibility or recombination. However, the genomic context, organization, and variation of these mating loci remain uncharacterized across Rhizoctonia fungi due to limited availability of well curated genome assemblies.

High-quality haplotype-aware genome assemblies have revealed substantial genomic divergence between nuclei coexisting within an individual strain of Rhizoctonia fungi. For example, in *Rhizoctonia cerealis*, two nuclear haplotypes can be differentiated by 427,567 SNPs as well as structural variants, duplications, inversions, and translocations (Han et al., 2025). Similar extensive divergence has also been identified and reported between haplotypes of *R. solani* AG8 (Han et al., 2025; Sperschneider et al., 2026). Importantly, heterozygosity is not limited to sequence divergence. In dikaryotic rust fungi, coexisting nuclear haplotypes can differ in gene content and expression, while somatic exchange of intact nuclei between lineages can generate new nuclear combinations associated with changes in host adaptation (Henningsen et al., 2025, 2024). A heterokaryon can therefore contain distinct nuclear genomes with substantial genetic divergence and potential to contribute differently to isolate biology and ecology, reflecting variation that cannot be fully represented by a single consensus genome. However, haplotype-resolved genomic comparisons remain limited, leaving the extent of nuclear divergence within *R. solani* AG1-IB and other related anastomosis groups unknown.

Here, we present to our knowledge the first haplotype-aware chromosome-scale genome assembly and heterokaryotic in *R. solani* AG1-IB strain S1W6. This strain was sampled from a tall fescue (*Lolium arundinaceum* (Schreb.) Darbysh.) brown patch disease lesion and has served as an experimental model in four research studies of pathogen coinfection, disease progression and plant-microbe interactions (Geyer et al., 2026; Green et al., 2024; Grunberg et al., 2023; O’Keeffe et al., 2021). The genome of strain S1W6 has provided a foundation for chromosome-level comparisons within AG1-IB and among other *R. solani* anastomosis groups (AGs), while enabling investigation of genomic organization and evolution of mating-type loci, an important but poorly understood aspect of Rhizoctonia fungi reproductive biology.

## 2. Materials and Methods

### 2.1 Fungal Isolation and Characterization

*Rhizoctonia solani* AG1-IB strain S1W6 was isolated in 2015 from tall fescue (*Lolium arundinaceum*) leaves exhibiting brown patch symptoms, collected from Duke Forest, North Carolina, USA (36.008 N, 79.018 W). Small sections of diseased tissue at the lesion margins were excised, surface disinfested in 2% NaOCl, rinsed in sterile water, and plated on Potato Dextrose Agar (PDA) (Becton, Dickinson and Company, Franklin Lakes, NJ) amended with chloramphenicol (400 mg/L). Emerging fungal hyphal tips were transferred to fresh PDA plates to obtain an axenic culture for nucleic acid extraction and sequencing.

### 2.2 DNA and RNA Extraction

S1W6 was cultured at 25 °C in Difco^TM^ Potato Dextrose Broth (PDB) (Becton, Dickinson and Company, Franklin Lakes, NJ) supplemented with ampicillin (100 mg/L). Mycelium was harvested through sterile cheesecloth and immediately pulverized in liquid nitrogen using a sterile mortar and pestle. The pulverized sample (∼500 mg) was aliquoted among 2.0 ml Eppendorf™ DNA LoBind™ Tubes (Eppendorf, Framingham, MA) and stored at −80 °C until use for DNA or RNA extraction, or Hi-C library preparation.

High molecular weight (HMW) DNA was extracted using a modified cetyltrimethylammonium bromide (CTAB) protocol adapted from Conlon et al., (2022). The modifications included: (i) treatment with RNase A at 37 °C for 30 min in OPS CTAB Extraction Buffer (OPS Diagnostics, Lebanon, NJ) prior to incubation with β-mercaptoethanol and proteinase K; (ii) precipitation of DNA using 0.7 volumes of ice-cold isopropanol relative to supernatant volume; and (iii) washing the DNA pellet three times with 70% ice-cold EtOH, followed by three washes with ice-cold absolute EtOH. DNA fragment length (>60 Kb) and integrity (DIN=8.9) were assessed using the Genomic DNA ScreenTape Analysis on the Agilent TapeStation (Agilent Technologies, Santa Clara, CA). Total RNA was extracted from frozen pulverized mycelium using QIAzol (QIAGEN Sciences Inc, Germantown, MD) according to the manufacturer’s instructions.

### 2.3 Library Preparation and Sequencing

High-molecular weight (HMW) genomic DNA (gDNA) libraries were prepared using the Native Barcode kit v14 and sequenced on a PromethION flow cell (Oxford Nanopore Technologies, Oxford, United Kingdom). Additional HMW gDNA libraries were prepared using the Rapid Barcode kit v14 and sequenced using MinION and PromethION platforms (Oxford Nanopore Technologies, Oxford, United Kingdom). Sequence output files were concatenated and combined for downstream analyses.

Dual-indexed Hi-C libraries to capture chromatin conformation and enable chromosome-scale scaffolding were prepared with the Proximo^TM^ Fungal Hi-C Kit (Phase Genomics, Seattle, WA) by following the manufacturer’s instructions (https://info.phasegenomics.com/hubfs/Protocol_Fungal_v4.5_202404.pdf) (Lieberman-Aiden et al., 2009; van Berkum et al., 2010). Libraries were sequenced on one lane of a NovaSeq X 10B flow cell (150 bp PE) (Illumina, Inc., San Diego, CA) at the High Throughput Sequencing Facility (HTSF) (The University of North Carolina at Chapel Hill, Chapel Hill, NC). Illumina sequencing libraries were generated for assembly polishing and quality control. Libraries were prepared with the Watchmaker DNA Library Prep Kit (Watchmaker Genomics) and sequenced together with the Hi-C libraries on one lane of a NovaSeq X 10B flow cell (150 bp PE) (Illumina, Inc., San Diego, CA) at the HTSF.

Poly-A RNA libraries were prepared using the NEBNext UltraExpress® RNA Library Prep Kit (New England Biolabs, Inc., Ipswich, MA) and sequenced on two lanes of a NovaSeq X 10B flow cell with 150 bp paired-end reads (Illumina, Inc., San Diego, CA) at the Genomic Sciences Laboratory (GSL) (North Carolina State University, Raleigh, NC) to support transcriptome-informed genome annotation.

### 2.4 Genome Assembly and Quality Control

*De novo* genome assembly was performed using PECAT v0.0.3 (Nie et al., 2024) on the DeCIFR platform (Carbone and White, 2025). PECAT assembly parameters were modified from default settings to increase stringency. The minimum read length was set to 5 kb, preprocessing and corrected-read coverage were set to 60x, and overlap filtering thresholds for alignment length and ratio, along with a minimum correction coverage of 6, were increased to reduce noise from short and repetitive reads (Nie et al., 2024). PECAT generated two sets of assemblies: (i) the haplotype 1 (Hap1) and haplotype 2 (Hap2) assemblies and (ii) the primary and alternate assemblies. The Hap1 and Hap2 assemblies were subsequently scaffolded using Hi-C data and used for all downstream analyses. The primary and alternate assemblies were also retained and made publicly available as additional genomic resources.

Hi-C reads were indexed and aligned using Chromap v0.3.2, followed by scaffolding with YaHS v1.2.2, while sequence indexing was performed using SAMtools v1.22.2 (Danecek et al., 2021; Zhang et al., 2021; Zhou et al., 2023). Hi-C data processing was performed using Juicer v1.6 (Durand et al., 2016). Final visualization of Hi-C contact maps was performed in Python using Cooler v0.10.4, NumPy v2.5.1, and Matplotlib v3.11.1 (Abdennur and Mirny, 2020; Harris et al., 2020; Hunter, 2007). Hi-C analyses were downloaded and executed using the Seqera Platform and Nextflow v24.10.5 build 5935 (Di Tommaso et al., 2017). Scaffolds were manually curated to correct misassemblies and improve chromosome-scale structure. After assembly, contigs shorter than 20 kb were removed. Small contigs corresponding to fragmented mitochondrial sequences produced during the PECAT assembly were identified, removed from the nuclear assembly, and replaced with a complete mitochondrial genome assembled separately through mitochondrial read enrichment and *de novo* assembly in Flye v2.9.6 (Kolmogorov et al., 2019). Mitochondrial reads were identified from Oxford Nanopore Technologies (ONT) whole-genome sequencing reads by mapping against the mitochondrial genome of *R. solani* AG1-IB, 7/3/14 accession HF546977.1 (Wibberg et al., 2013) using minimap2 v2.28 (Li, 2018). Raw reads with mitochondrial alignments were extracted using SeqKit v2.13.0 (Shen et al., 2016).

Due to excessive mitochondrial read depth causing assembly graph instability, the enriched mitochondrial read set was randomly downsampled to 5,000 reads using SeqKit. The downsampled read set was assembled using Flye v2.9.6 (Kolmogorov et al., 2019). Scaffolded nuclear and mitochondrial assemblies were polished using PolyPolish v0.6.1 (Wick and Holt, 2022), incorporating Illumina short reads to improve base-level accuracy. Reads were aligned to the assembly using BWA-MEM2 v2.3 (Vasimuddin et al., 2019) to generate a BAM file, which was subsequently used by PolyPolish for SNP-level error correction.

Raw read quality control was performed using fastp v0.23.2 (Chen et al., 2018) and MultiQC v1.11 (Ewels et al., 2020). Assembly completeness was assessed using BUSCO v6.0.0 (Simão et al., 2015) with the Basidiomycota odb12 database (Tegenfeldt et al., 2025), using miniprot v0.18-r281 (Li, 2023) for gene prediction and HMMER v3.4 (Eddy, 2011) for ortholog detection. Assembly quality metrics, including genome size, contiguity (N50), and GC content, were assessed using SeqKit v2.13.0 (Shen et al., 2016).

Fastp-filtered genomic short reads were independently mapped to the assemblies using BWA-MEM2 v2.3 to identify variant sites (Vasimuddin et al., 2019). For each mapping, nucleotide support from short reads across the genome was quantified using bcftools mpileup v1.10 at a maximum depth of 250 to calculate allelic depth and total read depth, which were subsequently used to calculate allelic read balance (Danecek et al., 2021). Analyses were restricted to biallelic SNPs called as heterozygous under the diploid model and having a minimum sample read depth of 10. For each SNP, the fraction of reads supporting the non-reference allele was calculated by dividing the number of reads supporting the non-reference allele by the total number of reads supporting either the reference or non-reference allele.

### 2.5 Comparative Genomics

Comparative genomic analyses were performed using FastANI v1.3 (Jain et al., 2018) to assess pairwise average nucleotide identity within and among AG1-IA, AG1-IB, and AG1-IC. This analysis included the newly generated S1W6 assembly alongside publicly available genomes within AG1. Metadata of strains included in the FastANI analysis are provided in **Table 1**. To facilitate comparison of assembly completeness, all previously published genomes were also assessed with BUSCO Basidiomycota odb12 **(Table 1)**. Chromosome numbering was assigned based on homology to the AG1-IA reference genome XN, as determined by whole-genome alignments using D-GENIES, rather than scaffold size. The Hap1 and Hap2 assemblies were restricted to chromosomes 1 through 16 and compared using MUMmer v4.0.0rc1, with alignments generated by nucmer and summarized using dnadiff to estimate the number of SNP differences between the haplotypes (Marçais et al., 2018). Genome-wide alignments of the Hap1 and Hap2 assemblies were conducted using D-GENIES v1.5.0 (Cabanettes and Klopp, 2018).

**Table 1.** *Rhizoctonia solani* AG1 genome assemblies included in comparative analyses and their BUSCO completeness metrics assessed using the Basidiomycota odb12 dataset.

| AG | Strain | Host Isolated From | Assembly accession | Origin | Source | BUSCO Complete (%) |
|---|---|---|---|---|---|---|
| AG1-IA | 1802/KB | *Oryza sativa* | GCA_001899475.2 | Kota Bharu, Malaysia | Mat Razali et al., 2021 | 83.6 |
| AG1-IA | YN-7 | *Oryza sativa* | GCA_015341985.1 | Yangzhou, China | Lee et al., 2021 | 95.5 |
| AG1-IA | B2 | *Oryza sativa* | GCA_015342405.1 | Arkansas, USA | Lee et al., 2021 | 95.1 |
| AG1-IA | ADB | *Oryza sativa* | GCA_015342415.1 | Adilabad, Telangana, India | Lee et al., 2021 | 94.4 |
| AG1-IA | WGL | *Oryza sativa* | GCA_015342435.1 | Warangal, Telangana, India | Lee et al., 2021 | 94.9 |
| AG1-IA | XN | *Zea mays* | GCA_016906535.1 | China | Li et al., 2021 | 95.2 |
| AG1-IA | ZJ | *Setaria italica* | GCA_019297585.1 | Zhejiang, China | Liu et al., 2024 | 94.6 |
| AG1-IA | HG81 | *Oryza sativa* | GCA_024722335.1 | Hubei, China | Yang et al., 2022 | 96.2 |
| AG1-IA | BRS1 | *Oryza sativa* | GCA_028551635.1 | India | Francis et al., 2023 | 95.3 |
| AG1-IA | RAP2 | *Echinochloa frumentacea* | GCA_044039995.1 | Andhra Pradesh, India | Patro et al., 2024 | 94 |
| AG1-IA | D138 | Unknown | GCA_045866225.1 | Unknown | GenBank Assembly | 95.9 |
| AG1-IA | BM1 | *Glycine max* | GCA_905219565.1 | Unknown | Kaushik et al., 2022 | 92.4 |
| AG1-IA | VAP1 | *Panicum miliaceum* | GCA_048471615.1 | Andhra Pradesh, India | Koti et al., 2025 | 91.3 |
| AG1-IB | 7/3/14 | *Lactuca sativa* | GCA_000832345.2 | Großbeeren, Germany | Wibberg et al., 2015 | 95.2 |
| AG1-IB | LK4 | *Nicotiana tabacum* | GCA_026766895.1 | Guizhou, China | Unpublished | 95.7 |
| AG1-IB | BM2 | *Lolium arundinaceum* | GCA_905219545.1 | Unknown | Kaushik et al., 2022 | 95.8 |
| AG1-IB | O8/2 | *Lactuca sativa* | GCA_914271495.1 | Germany | Wibberg et al., 2021 | 95.4 |
| AG1-IB | 1/2/21 | *Lactuca sativa* | GCA_914271525.1 | Germany | Wibberg et al., 2021 | 94.6 |
| AG1-IB | S1W6 (Hap1) | *Lolium arundinaceum* | JCBRKY000000000 | NC, USA | *This study* | 96.2 |
| AG1-IB | S1W6 (Hap2) | *Lolium arundinaceum* | JCBRKZ000000000 | NC, USA | *This study* | 96.1 |
| AG1-IC | BM3 | *Beta vulgaris* | GCA_905219585.1 | Unknown | Kaushik et al., 2022 | 96.2 |

### 2.6 Genomic Characterization

RNA-seq reads were quality filtered using fastp v0.23.2 (Chen et al., 2018) and summarized with MultiQC v1.11 (Ewels et al., 2020), then assembled *de novo* using Trinity v2.15.2 (Grabherr et al., 2011) to generate transcript evidence for genome annotation. Genome annotation was performed using MAKER v2.31.11 (Cantarel et al., 2008; Holt and Yandell, 2011), integrating Trinity-assembled transcripts with *ab initio* gene predictions from AUGUSTUS v3.2.3 (Stanke et al., 2008; Stanke and Waack, 2003). A custom AUGUSTUS model was trained using gene models from a closely related species *R. solani* AG1-IA isolate XN (Li et al., 2021) and incorporated into the MAKER pipeline. Both assemblies were annotated using the same approach.

Annotated protein sequences were obtained from MAKER and input into the online graphical user interface of eggNOG-mapper v3.0.0-beta6 for proteins encoded on Chromosome 1 through 16. Functional annotation was performed with eggNOG-mapper v3.0.0-beta6 (Cantalapiedra et al., 2021) using orthology assignments from the eggNOG 7 database (Hernández-Plaza et al., 2026). Homology searches were carried out with DIAMOND in sensitive mode with iterative search enabled (ortholog e-value ≤ 0.001). Gene Ontology (GO) were transferred from the identified orthologs within the automatically adjusted taxonomic scope (--tax_scope auto). GO identifiers were mapped to term names and ontology categories, biological process, cellular component, and molecular function, using the GO.db v3.21.0 (Carlson, 2026) and AnnotationDbi v1.70.0 packages in R (Pagès et al., 2025). To assess differences in GO term usage between assemblies, gene counts per GO term were compared using Fisher’s exact tests. Resulting p-values were corrected for multiple testing using the Benjamini-Hochberg procedure to control the false discovery rate (FDR) and GO terms with adjusted p-values less than 0.05 were considered significantly different.

Homeodomain (HD) proteins and *STE3* were identified by BLAST searches performed in Geneious Prime version 2026.1 (Biomatters Ltd.) using reference HD genes from *R. solani* AG1-IA XN and *Ceratobasidium* AG-Ba JN and LY (Camacho et al., 2009; Kearse et al., 2012; Li et al., 2021). Identified HD genes were translated into the amino acid sequences and aligned in MAFFT v7.526 (Katoh and Standley, 2013). To explore the presence and absence of HD and *STE3* homologs across AG1, all genome assemblies included in this study were searched using TBLASTN v2.16.0+ with HD and STE3 protein sequences from XN and S1W6 as queries. HD and *STE3* homologs were considered present when at least one corresponding query produced a hit with an E-value less than 10⁻⁵⁰ and greater than 50% query coverage. Predicted homeodomain protein sequences were analyzed using SUPERFAMILY 2.0 to identify conserved protein domains (Pandurangan et al., 2019).

Putative effectors were identified using the Predector pipeline (Jones et al., 2021), installed and executed using the Seqera Platform and Nextflow v24.10.5 build 5935 (Di Tommaso et al., 2017). This pipeline incorporates SignalP 6.0 (Teufel et al., 2022), EffectorP 3.0 (Sperschneider and Dodds, 2022), and TMHMM 2.0 (Krogh et al., 2001). SignalP was used to identify proteins containing predicted signal peptides indicative of secretion. These secreted proteins were classified using EffectorP, which predicts putative effectors and assigns likely localization as cytoplasmic, apoplastic or dual. Finally, TMHMM was used to exclude proteins containing predicted transmembrane helices from the effector set. TMHMM was also used on the translated sequences of *STE3*-like genes in S1W6 and XN AG1-IA reference strain (Krogh et al., 2001). Carbohydrate-active enzymes (CAZymes) were annotated using the Predector pipeline, which leverages dbCAN3 version 3.3.2 to identify CAZyme content and classify proteins into functional families (Zheng et al., 2023). Biosynthetic gene clusters (BGCs) were identified using the fungal version of antiSMASH v8.0.4 with strict detection settings (Blin et al., 2025).

## 3. Results

### 3.1 Genome Sequencing, Assembly, and Quality Control

The PECAT assembly produced haplotype 1 (Hap1) and haplotype 2 (Hap2) assemblies. PECAT separates homologous sequence based on haplotype variation in the assembly graph but does not assign the resulting chromosome homologs to defined nuclear backgrounds. *R. solani* strain S1W6 was isolated as a heterokaryon, but the identity, genomic composition and origin of the monokaryotic (haploid) lineages are unknown. Consequently, the reconstructed chromosome homologs cannot be assigned to independently defined nuclear lineages, and the PECAT designations Hap1 and Hap2 are retained throughout this study. The final Hap1 assembly had a total length of 41.7 Mb (N50 = 2.48 Mb; GC = 48.2%), while the Hap2 assembly had a total length of 39.4 Mb (N50 = 2.30 Mb; GC = 48.2%). Genome completeness, assessed using BUSCO, was 96.2% for Hap1 and 96.1% for Hap2, with 0.5% duplicated BUSCOs in both assemblies and 0.6% and 0.7% fragmented BUSCOs, respectively **(Table 1)**. A total of 11,987 and 11,483 genes were predicted in the Hap1 and Hap2 assemblies, respectively. The mitochondrial genome was assembled as a single circular contig of 173,371 bp with a mean Oxford Nanopore sequencing coverage of 397x. PolyPolish indicated high assembly accuracy, with estimated pre-polishing accuracies ranging from 99.97 to over 99.99%. All scaffolds had mean coverage exceeding 1000x; additional polishing statistics are provided in **Supplementary Table 1**. Hi-C contact maps revealed distinct chromatin interaction patterns between the haplotypes **(Supplementary Figure 1).** Approximately 72% of heterozygous biallelic SNPs exhibited balanced allele frequencies ranging from 0.40 to 0.60 in both assembly mappings, consistent with approximately equal representation of the two nuclear haplotypes in the sequencing data.

### 3.2 Comparative Genomics

No major structural rearrangements were detected between the assemblies aligned in D-GENIES (**Supplementary Figure 2**). Pairwise comparison of haplotypes using MUMmer alignments and dnadiff indicated that S1W6 Hap1 and Hap2 assemblies differed by 646,037 SNPs and 88,990 indels with 97.75% average alignment identity. Pairwise Average Nucleotide Identity (ANI) analysis using FastANI showed distinct separation of AG1-IA, AG1-IB and AG1-IC (**Figure 1**). Genomes within AG1-IA exhibited high similarity, with ANI values greater than 96% with some substructuring and clustering, whereas comparisons between AG1-IA and AG1-IB or AG1-IC ranged from 80 to 82%, consistent with strong divergence between subgroups. Within AG1-IB, S1W6 clustered with previously published assemblies 7/3/14 (GCA_000832345.2) and BM2 (GCA_905219545.1), sharing 97.2 to 97.8% of ANI values with other AG1-IB genomes. The S1W6 Hap1 and Hap2 assemblies exhibited 97.5% ANI when measured by FastANI, corresponding to approximately 2.5% divergence between haplotypes residing within the same cells, a level of variation comparable to that observed between unrelated AG1-IB isolates from different hosts located different countries.

**Figure 1.**
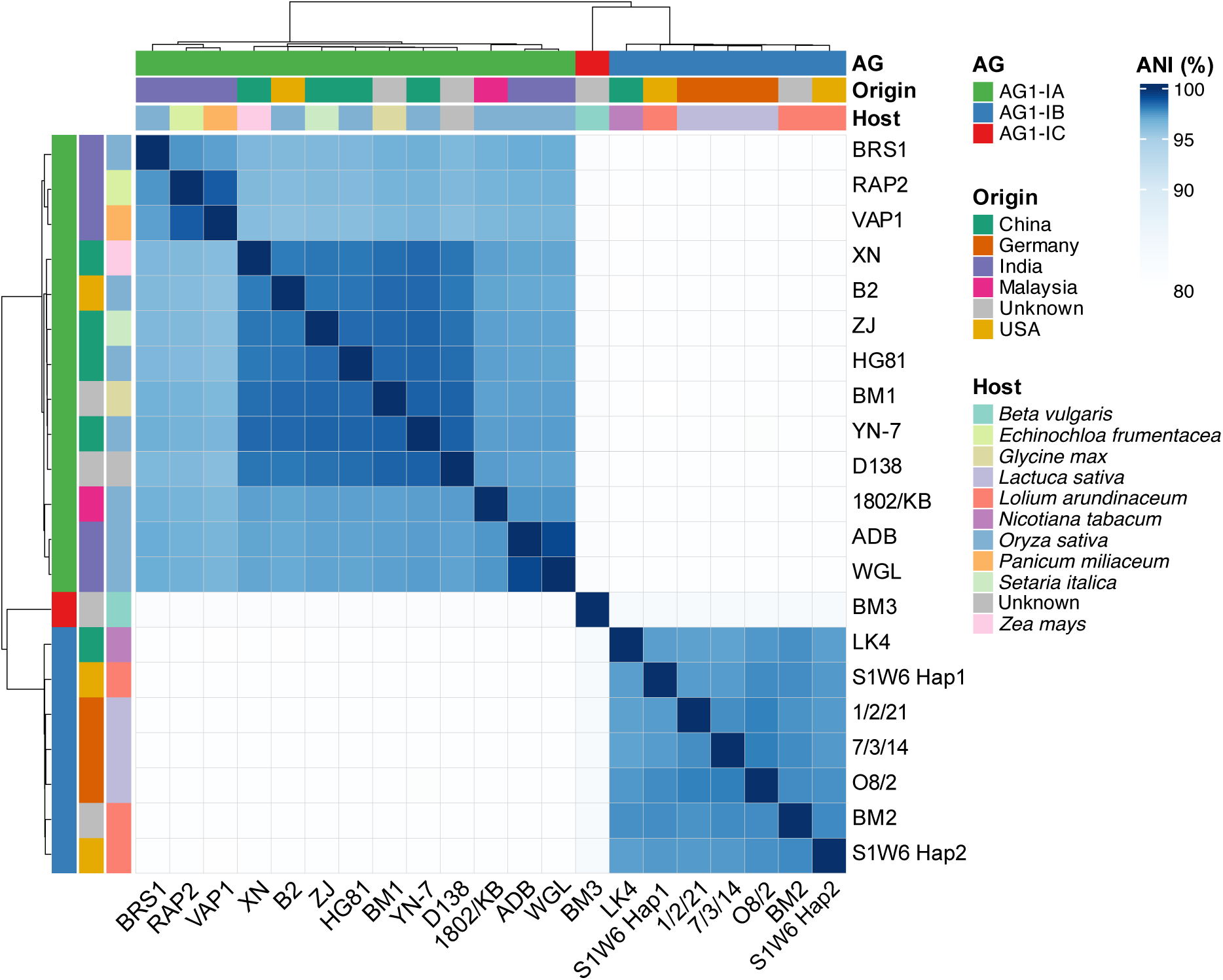
Pairwise average nucleotide identity (ANI) among publicly available *Rhizoctonia solani* AG1 genomes and the S1W6 Hap1 and Hap2 assemblies. Color annotations indicate anastomosis group (AG), geographic origin, and host species.

### 3.3 Genomic Characterization

The HD genes were located on chromosome 12 in both assemblies. Hap2 contained four HD genes (*W-HD1*, *W-HD2*, *E-HD1*, *E-HD2*), whereas Hap1 lacked *E-HD1*. HD genes shared between assemblies (*W-HD1*, *W-HD2*, *E-HD2*) showed sequence variation (**Figure 2**). All predicted HD proteins contained conserved homeodomain DNA-binding domains identified through the SUPERFAMILY database and are presented in **Supplementary Table 2**. The identified HD genes were translated into amino acid sequences and aligned by homolog across the S1W6 assemblies and the reference assembly, isolate XN. Comparative visualization of the HD locus revealed differences in organization and HD sequence conservation among assemblies. E-HD2 proteins were highly divergent: the S1W6 copies shared 63.2% amino acid identity, while the Hap1 E-HD2 allele was more similar to XN (80.8%) than was the Hap1 E-HD2 allele (60.8%) **(Figure 2a)**. In contrast, W-HD1 proteins were comparatively conserved, with the S1W6 Hap1 and Hap2 copies sharing 90.5% amino acid identity; both copies of W-HD1 encoded in S1W6 were also more similar to each other than to XN, retaining 77.7% and 76.2% identity for the Hap1 and Hap2 alleles. W-HD2 proteins exhibited substantially greater divergence, with 61.6% identity between the S1W6 Hap1 and Hap2 copies and only 47.2% and 43.1% identity between XN and the S1W6 Hap1 and Hap2 copies, respectively. Polymorphic amino acid sites within W-HD1, W-HD2, and E-HD2 were concentrated primarily within the N- and C-terminal regions of the proteins **(Figure 2b)**. The homeodomain-like regions predicted by SUPERFAMILY remained largely conserved and lacked detectable amino acid polymorphisms between the alternative alleles within S1W6.

**Figure 2.**
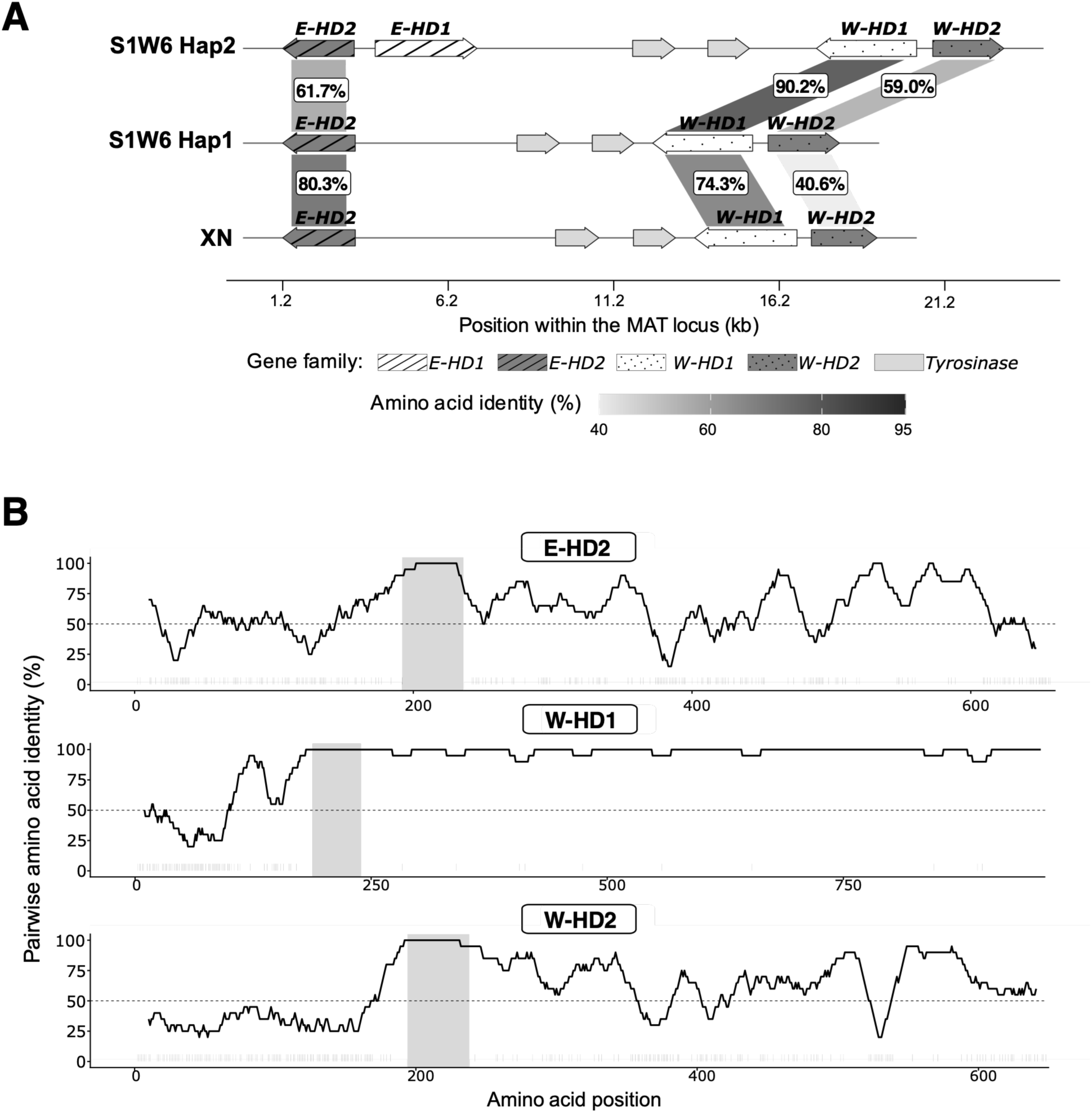
**(A)** Syntenic comparison of the homeodomain (HD) mating-type locus among the S1W6 Hap1 and Hap2 assemblies and *R. solani* AG1-IA isolate XN. Pairwise amino acid identities between homologous HD proteins are shown, with shaded ribbons indicating the degree of amino acid identity. **(B)** Sliding-window profiles (20 amino acids per window) of pairwise amino acid identity for E-HD2, W-HD1, and W-HD2 between the haplotypes of S1W6. Tick marks indicate variable amino acid positions, and gray shaded regions denote predicted homeodomain-like regions identified using SUPERFAMILY.

A *STE3* gene homologous to fungal pheromone receptor genes involved in mating was identified on chromosome 15. Unlike HD proteins, STE3 proteins in S1W6 were highly conserved, sharing 99.7% amino acid identity and differing at a single amino acid position **(Figure 3)**. The STE3 proteins from S1W6 Hap1 and Hap2 showed high amino acid sequence identity to XN (94.0% and 94.3%, respectively). TMHMM predicted six of the seven expected transmembrane domains characteristic of fungal STE3 pheromone receptors in S1W6 and XN (Xue et al., 2008). Although seven peaks in posterior transmembrane probability were observed, one peak did not meet the confidence threshold for classification as a discrete transmembrane helix. Posterior probability plots and predicted transmembrane domains from TMHMM are presented in **Supplementary Figure 3**. Using S1W6 and XN protein sequences as queries, TBLASTN searches detected at least one high-confidence HD homolog and one STE3 homolog in each of the 21 AG1 genome assemblies examined, with qualifying hits covering at least half of the query sequence and having E-values below 10^-50^. Given the variable quality of available assemblies, including collapsed or consensus representations that may obscure haplotype-specific variation, the broader survey was limited to homolog detection, while detailed characterization of mating-type loci was restricted to S1W6 and XN. Gene Ontology (GO) annotation profiles were consistent between the assemblies when restricted to chromosomes Chr01 through Chr16. This analysis included 10,593 genes from Hap1 and 11,083 genes from Hap2, with no significant differences among the 11,973 GO terms tested (minimum unadjusted p = 0.190; all FDR-adjusted p-values were greater than 0.05). Annotated gene models were further analyzed to identify putative effectors. The Hap1 assembly contained 87 cytoplasmic, 253 apoplastic, and 65 dual-localized effectors (405 total), while the Hap2 assembly contained 84 cytoplasmic, 255 apoplastic, and 62 dual-localized effectors (401 total) **(Supplementary Table 3)**. CAZyme class composition was similar between haplotypes, with 1,152 unique CAZyme proteins predicted in Hap1 and 1,125 in Hap2 **(Supplementary Table 3).** Biosynthetic gene clusters (BGCs) were identified using fungiSMASH v8.0.4 (Blin et al., 2025). BGC composition was similar between haplotypes, with 12 clusters in Hap1 and 13 in Hap2, including two nonribosomal peptide synthetase (NRPS) clusters in each **(Supplementary Table 4).** Hap2 contained an additional predicted terpene cluster on Chr06, bringing the total number of terpene clusters to 11 compared with 10 in Hap1.

**Figure 3.**
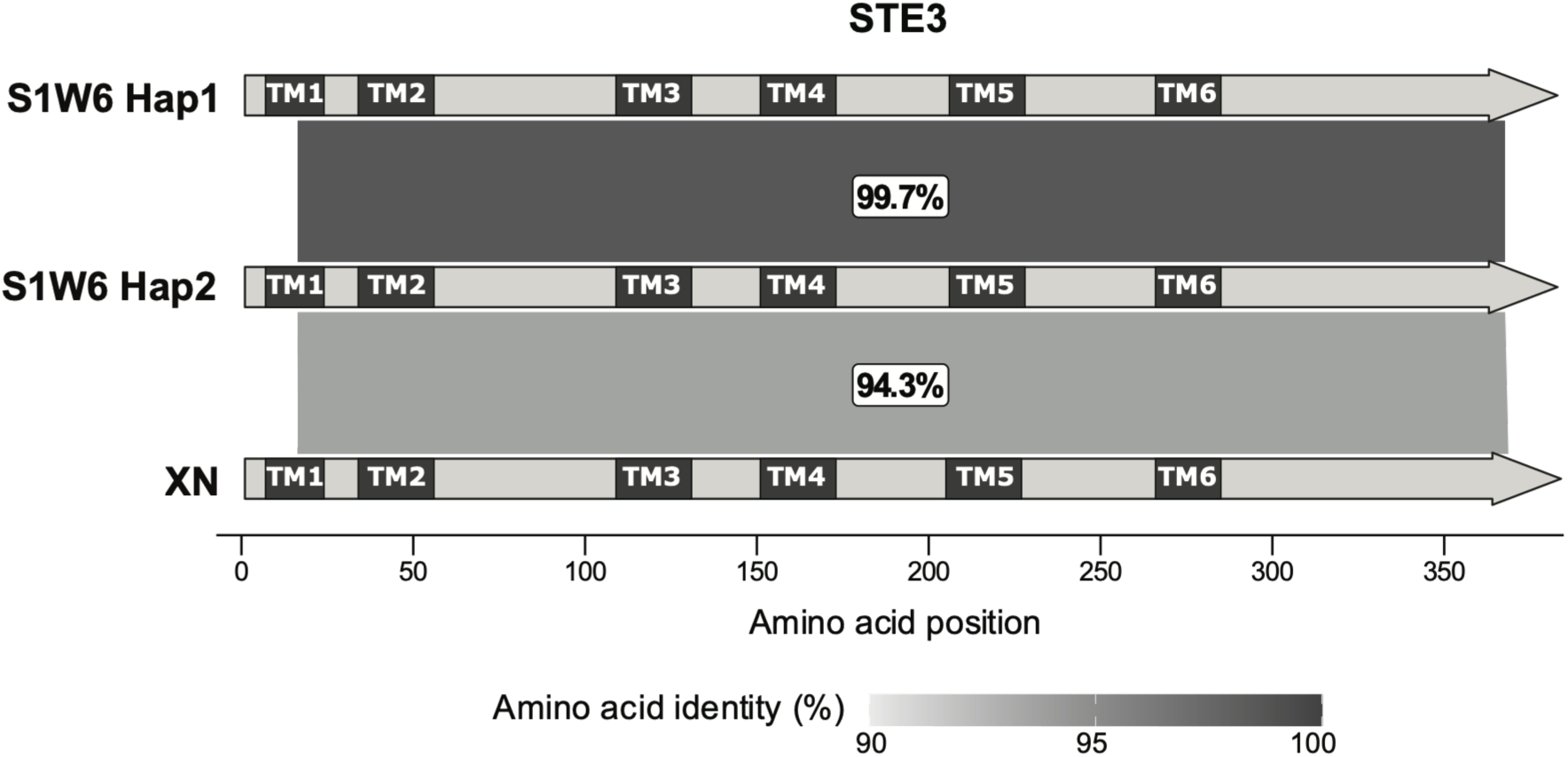
Comparison of STE3 pheromone receptor protein architecture and pairwise amino acid identity among S1W6 Hap1 and Hap2 and *R. solani* AG1-IA isolate XN. Predicted transmembrane domains (TM1-TM6) are indicated for each STE3 homolog. Shaded ribbons indicate the degree of amino acid identity between homologs.

## 4. Discussion

Existing genome assemblies of *Rhizoctonia solani* AG1-IB have relied exclusively on short-read sequencing, resulting in fragmented draft genomes that lack haplotype resolution. Although genome assembly continuity has improved over time, in general, most publicly available AG1-IB genomes provide incomplete chromosome scale resolution, limiting analyses of chromosome organization, structural variation, and heterokaryotic genomic architecture. Consequently, fundamental questions regarding genome evolution, nuclear diversity and pathogenicity in this multinucleate fungus have remained challenging to address. This study provides the first haplotype-resolved, chromosome-scale assembly of a multinucleate AG1-IB heterokaryon and enabled the first comprehensive characterization of the candidate HD and *STE3* mating-type loci. Our study supported the conservation of the *STE3* pheromone receptor gene but revealed extensive divergence among homeodomain (HD) genes.

The pheromone receptor gene *STE3*, a member of the fungal seven-transmembrane pheromone receptor family, was identified on chromosome 15, and the two copies detected in S1W6 were highly conserved (Xue et al., 2008). TMHMM detected six transmembrane domains with high confidence in the *STE3* amino acid sequences recovered from the Hap1 and Hap2 assemblies of S1W6 and in the AG1-IA reference strain XN. While the six domains had high posterior probabilities supporting the presence of transmembrane domains, there was one additional putative transmembrane domain with a low posterior probability not predictive of a transmembrane domain function (**Supplementary Figure 3**). The S1W6 STE3 protein sequences were 99.7% identical, with a single amino acid residue varying between assemblies at position 159: isoleucine in the Hap1 assembly and leucine in the Hap2 assembly. The strong conservation of the amino acid sequence of STE3 contrasts sharply with the divergence observed between homeodomain mating genes, particularly E-HD2 (61.7% amino acid identity) and W-HD2 (59.0% amino acid identity) between haplotypes. HD and STE3 homologs were broadly detected across available AG1 genomes; however, most were generated as collapsed or consensus assemblies from heterokaryotic isolates, limiting reliable comparisons of mating-locus gene content and organization.

Although the overall functional gene repertoire was highly conserved between assemblies, localized divergence was evident at the HD mating locus. The Hap1 assembly contained *W-HD1, W-HD2, and E-HD2,* mirroring the organization reported in AG1-IA strain XN **(Figure 2)**, used as a reference and control strain in this study, whereas the Hap2 assembly additionally contained *E-HD1*. As in XN, we identified two putative tyrosinases between the *W-HD1*/*W-HD2* and *E-HD1*/*E-HD2* gene pairs, suggesting that tyrosinases may be a consistent feature of the *R. solani* HD locus. Further, each HD1/HD2 gene pair was divergently transcribed when both copies were present, consistent with model HD fungal mating loci in *Coprinopsis cinerea* (Pardo et al., 1996). All HD gene amino acid sequences were classified as containing homeodomains, consistent with their expected functional role. In Basidiomycota fungi, compatible HD1 and HD2 proteins heterodimerize to form transcription factors, the formation of which is facilitated by protein sequence variation (Kües, 2015; Spit et al., 1998). The functional HD heterodimer enters the nucleus and regulates genes required for dikaryotic development and sexual reproduction (Kües, 2015). In the model Basidiomycota mushroom-forming fungus *Coprinopsis cinerea* and wood decay fungus *Schizophyllum commune*, activation of the HD pathway coordinates nuclear pairing and division during establishment and maintenance of the dikaryon (Kües, 2015). In these fungi, HD signaling also regulates clamp-cell formation and contributes to the initiation of fruiting-body development (Kües, 2015). Interestingly, species of Rhizoctonia fungi, including *R. solani* and *Ceratobasidium*, lack clamp connections despite forming persistent heterokaryons and undergoing sexual reproduction (Cubeta and Vilgalys, 1997), leaving the developmental functions regulated by compatible HD1-HD2 heterodimers in Rhizoctonia fungi unresolved.

Expression of all predicted HD genes under laboratory culture conditions indicates that these loci are transcriptionally active and represent strong candidates for functional characterization as mating determinants. The identified HD proteins exhibited substantial sequence divergence between the S1W6 assemblies, although the degree of divergence differed among HD gene classes. E-HD2 and W-HD2 protein sequences were particularly variable, with only 61.7% and 59.0% amino acid identity between the Hap1 and Hap2 copies, whereas W-HD1 remained comparatively conserved at 90.2% amino acid identity. The S1W6 Hap1 E-HD2 allele shared greater amino acid identity when aligned with the AG1-IA strain XN (80.3%) than with the corresponding S1W6 Hap2 E-HD2 allele (61.7%). Although this observation is based on a limited number of protein sequences, it suggests that trans-species polymorphism may occur in HD genes across *R. solani*, a hypothesis that warrants further investigation. In Basidiomycota fungal mating systems, heterodimerization specificity between HD proteins is mediated primarily through the N-terminal interaction domains, while the homeodomain region remains comparatively conserved (Banham et al., 1995; Spit et al., 1998). Consistent with this pattern, the predicted homeodomain-like regions of W-HD1, W-HD2, and E-HD2 lacked amino acid polymorphisms within *R. solani* strain S1W6 despite broader sequence divergence across these proteins.

Despite extensive sequence divergence at the HD locus, genome-wide functional content remained largely conserved. GO annotation profiles did not differ significantly between the Hap1 and Hap2 assemblies, suggesting that substantial nucleotide sequence divergence can accumulate between coexisting nuclei while preserving much of the broader functional gene repertoire.

The increased level of resolution gained by using a haplotype aware method enables explicit representation of haplotypic variation within a heterokaryon, providing a more complete model than previously reported in AG1. In *R. cerealis*, which possesses two nuclei per hyphal cell, haplotype-resolved assemblies revealed extensive within-isolate divergence, including over 400,000 SNPs and substantial structural variation between nuclear backgrounds (Han et al., 2025). Similarly, haplotypes of *R. solani* AG8 differed by 551,138-880,333 SNPs and exhibited 96.8-98.1% average alignment identity (Sperschneider et al., 2026). Consistent with these studies, pairwise comparison of the S1W6 Hap1 and Hap2 assemblies identified 646,037 SNPs, 88,990 indels, and an average alignment identity of 97.75%. Haplotype comparisons in S1W6 demonstrate that substantial within-isolate genetic variation is present in AG1-IB. Future comparative analyses across additional anastomosis groups are needed to determine whether highly heterozygous genomes are a common feature of Rhizoctonia fungi heterokaryons and how this diversity contributes to genome evolution and population diversity and structure.

The extent of within-isolate genetic variation observed in S1W6 approaches the variation observed among independently collected AG1-IB isolates, despite the two haplotypes coexisting within the same vegetative mycelium. Pairwise average nucleotide identity analysis showed that the S1W6 assemblies share 97.5% nucleotide identity across aligned regions, corresponding to approximately 2.5% divergence between haplotypes. This value was comparable to, and in some cases lower than, the ANI observed between S1W6 and other AG1-IB genomes, including isolates recovered from lettuce in Germany (7/3/14) and tobacco in China (LK4). Thus, genetic backgrounds coexisting within a single strain are no more similar to one another than they are to AG1-IB isolates collected from different plant hosts and continents. This finding suggests that heterokaryotic individuals can harbor levels of genetic variation approaching those observed across the broader AG1-IB population and highlights the potential for heterokaryosis to act as a reservoir of evolutionary potential and adaptability within a single biological individual.

Interestingly, FastANI analyses revealed evidence of genomic substructure within AG1-IA that was not apparent among the currently available AG1-IB genomes. Several AG1-IA isolates formed higher-identity clusters, in some cases among isolates recovered from the same host, suggesting that host-associated or population-level structure may be present within AG1-IA. In contrast, AG1-IB isolates recovered from lettuce in Germany (7/3/14, O8/2, and 1/2/21), tobacco in China (LK4), and tall fescue in North America (S1W6) and from tall fescue in an unreported location (BM2) exhibited broadly similar ANI values and limited evidence of host or geography-associated clustering. These patterns should be interpreted cautiously because many currently available AG1 genome assemblies are fragmented or represent consensus sequences that do not resolve the constituent haplotypes of heterokaryotic isolates. Consequently, pairwise ANI calculated from these assemblies may obscure within-isolate variation and relationships among haplotypes. Additional AG1-IB genome assemblies will be required to rigorously evaluate population structure within AG1-IB.

In contrast to the extensive genomic divergence between the S1W6 haplotypes, their predicted BGC repertoires were highly similar. The Hap1 assembly contained 10 terpene-associated clusters and two NRPS clusters, with the Hap2 assembly containing an additional terpene-associated BGC on chromosome 6. This pattern is consistent with observations in *R. solani* AG3 strains, where BGC repertoires are similarly dominated by terpene and NRPS/NRPS-like clusters (Xu et al., 2024). Likewise, previously analyzed AG1-IB genomes, 1/2/21, O8/2 and 7/3/14, contain 15 to 17 predicted secondary metabolite regions, including six NRPS clusters and multiple terpene-associated clusters (Wibberg et al., 2021). Overall, these findings suggest that BGC repertoires remain stable across AG1-IB despite substantial genomic divergence. Consistent with this broader pattern, predicted effector and CAZyme class distributions differed very little between haplotypes. However, this study did not evaluate sequence-level variation among effectors and CAZymes shared between haplotypes. Consequently, the extent of allelic diversity within these pathogenicity drivers, and the potential contribution to virulence or host adaptation, remains unknown in Rhizoctonia fungi.

The application of haplotype-aware assembly approaches represents a substantial advance for studying heterokaryotic fungi. By modeling both coexisting haplotypes within heterokaryotic individuals, these methods reveal within-strain variation, structural differences, and locus-specific divergence obscured in consensus assemblies. In S1W6, this approach uncovered extensive divergence at the candidate HD mating locus while demonstrating that much of the broader functional gene repertoire remains conserved between haplotypes. The deep divergence among HD alleles is consistent with the long-term maintenance of mating-type diversity expected under balancing selection, although additional functional studies are required to determine the roles of these loci in AG1-IB. Expanding genomic resources across these closely related subgroups that have the inherent capability of undergoing vegetative interactions involving hyphal fusion, nuclear and cytoplasmic exchange, will be important for understanding genetic diversity across AG1. Such efforts are critical given the broad host range and economic importance of *R. solani* as a plant pathogen (Dean et al., 2012). Improved understanding of the genomic variation underlying reproduction and host association may ultimately help inform disease management across diverse plant pathosystems. Our findings demonstrate the value of this approach by resolving substantial haplotypic and mating-locus variation within a single AG1-IB isolate. To our knowledge, S1W6 represents the most contiguous and complete heterokaryotic genomic resource currently available for AG1, providing a valuable reference for future investigations of heterokaryosis, mating systems, and genome evolution in *R. solani* and related *Rhizoctonia* fungi.

## CRediT authorship contribution statement

J.B.C.: Conceptualization, Methodology, Formal analysis, Software, Investigation, Visualization, Writing - original draft, Writing - review and editing. M.A.C.: Investigation, Methodology, Writing - review and editing. M.A.Cu.: Resources, Writing - review and editing. J.B.W.: Software. R.S.: Software. I.C.: Methodology, Resources, Supervision, Funding acquisition, Writing - review and editing, Writing - original draft.

## Data availability

The chromosome-scale annotated genome assemblies of *Rhizoctonia solani* AG1-IB isolate S1W6 are available through NCBI under BioSample accession SAMN62087558. Raw Oxford Nanopore PromethION and MinION genomic DNA reads, Illumina cDNA reads, Illumina whole-genome sequencing reads, and Illumina Hi-C reads are available through the NCBI Sequence Read Archive (SRA) under the same BioSample accession. Supplementary figures, tables, and data are available with the online version of this article. Genome annotations and analysis outputs from Predector, antiSMASH, eggNOG-mapper, and dnadiff are available in Dryad (https://doi.org/10.5061/dryad.4f4qrfjvt). The authors confirm that all data, code, and protocols supporting the findings of this study are provided within the article, its supplementary materials, or the associated public repositories.

## Acknowledgements

We thank Dr. Kayleigh O’Keeffe for collecting and isolating S1W6 and Dr. Charles Mitchell for assistance with securing grant funding. We are grateful to Jash Gopani for supporting the deployment of the Seqera Platform at North Carolina State University, enhancing the scalability and reproducibility of our genomic analyses.

