## Supplementary Information for "Chromosome-scale genome assembly of multinucleate heterokaryotic *Rhizoctonia solani* AG1-IB"

**Supplementary Table 1.** Polypolish SNP error correction and quality control statistics for the S1W6 Hap1 and Hap2 assemblies. Statistics include scaffold length, mean short-read depth, percentage of bases covered by short reads, and the number and percentage of nucleotide positions corrected during polishing. Estimated pre-polishing accuracy and Phred-scaled quality values (QV) were calculated from the proportion of nucleotide positions corrected.

| Assembly | Scaffold | Length (Mb) | Mean depth (×) | Coverage (%) | Positions changed | Changed (%) | Est. pre-polish accuracy (%) | QV |
|---|---|---|---|---|---|---|---|---|
| Hap1 | Chr01 | 3.756 | 1824.9 | 99.9646 | 46 | 0.0012 | 99.9988 | Q49.12 |
| Hap1 | Chr02 | 2.526 | 1884.2 | 99.9182 | 60 | 0.0024 | 99.9976 | Q46.24 |
| Hap1 | Chr03 | 3.690 | 2113.7 | 99.9659 | 81 | 0.0022 | 99.9978 | Q46.58 |
| Hap1 | Chr04 | 3.754 | 1903.4 | 99.8321 | 31 | 0.0008 | 99.9992 | Q50.83 |
| Hap1 | Chr05 | 2.326 | 1784.3 | 99.7637 | 38 | 0.0016 | 99.9984 | Q47.87 |
| Hap1 | Chr06 | 3.391 | 1699.0 | 99.9041 | 184 | 0.0054 | 99.9946 | Q42.65 |
| Hap1 | Chr07 | 2.225 | 1831.4 | 99.9134 | 50 | 0.0022 | 99.9978 | Q46.48 |
| Hap1 | Chr08 | 2.477 | 1767.6 | 99.9633 | 85 | 0.0034 | 99.9966 | Q44.65 |
| Hap1 | Chr09 | 2.141 | 1641.8 | 99.9030 | 26 | 0.0012 | 99.9988 | Q49.16 |
| Hap1 | Chr10 | 1.970 | 1737.0 | 99.8773 | 71 | 0.0036 | 99.9964 | Q44.43 |
| Hap1 | Chr11 | 2.241 | 1710.0 | 99.8526 | 36 | 0.0016 | 99.9984 | Q47.94 |
| Hap1 | Chr12 | 1.905 | 1734.3 | 99.9499 | 45 | 0.0024 | 99.9976 | Q46.27 |
| Hap1 | Chr13 | 2.080 | 1746.9 | 99.9104 | 46 | 0.0022 | 99.9978 | Q46.55 |
| Hap1 | Chr14 | 2.167 | 1813.3 | 99.9835 | 54 | 0.0025 | 99.9975 | Q46.03 |
| Hap1 | Chr15 | 1.661 | 1695.5 | 99.8852 | 41 | 0.0025 | 99.9975 | Q46.07 |
| Hap1 | Chr16 | 3.414 | 1904.7 | 99.8258 | 64 | 0.0019 | 99.9981 | Q47.27 |
| Hap1 | Mito | 0.173 | 25833.7 | 99.9988 | 0 | 0 | 100 | Q∞ |
| Hap2 | Chr01 | 3.599 | 1872.0 | 99.7361 | 94 | 0.0026 | 99.9974 | Q45.83 |
| Hap2 | Chr02 | 2.523 | 1901.1 | 99.9085 | 36 | 0.0014 | 99.9986 | Q48.46 |
| Hap2 | Chr03 | 3.239 | 2253.0 | 99.7881 | 102 | 0.0031 | 99.9969 | Q45.02 |
| Hap2 | Chr04 | 3.629 | 1916.0 | 99.8748 | 43 | 0.0012 | 99.9988 | Q49.26 |
| Hap2 | Chr05 | 2.302 | 1827.0 | 99.8881 | 47 | 0.0020 | 99.9980 | Q46.90 |

| | | | | | | | | |
|---|---|---|---|---|---|---|---|---|
| Hap2 | Chr06 | 3.146 | 1766.9 | 99.8857 | 109 | 0.0035 | 99.9965 | Q44.60 |
| Hap2 | Chr07 | 2.069 | 1896.4 | 99.9739 | 38 | 0.0018 | 99.9982 | Q47.36 |
| Hap2 | Chr08 | 2.026 | 1889.4 | 99.9496 | 45 | 0.0022 | 99.9978 | Q46.53 |
| Hap2 | Chr09 | 2.085 | 1761.0 | 99.8596 | 47 | 0.0023 | 99.9977 | Q46.47 |
| Hap2 | Chr10 | 1.956 | 1759.5 | 99.9426 | 44 | 0.0022 | 99.9978 | Q46.48 |
| Hap2 | Chr11 | 2.040 | 1743.7 | 99.9507 | 33 | 0.0016 | 99.9984 | Q47.91 |
| Hap2 | Chr12 | 1.876 | 1768.6 | 99.9368 | 35 | 0.0019 | 99.9981 | Q47.29 |
| Hap2 | Chr13 | 1.961 | 1787.7 | 99.9765 | 38 | 0.0019 | 99.9981 | Q47.13 |
| Hap2 | Chr14 | 1.982 | 1860.1 | 99.8906 | 38 | 0.0019 | 99.9981 | Q47.17 |
| Hap2 | Chr15 | 1.328 | 1876.3 | 99.8570 | 26 | 0.0020 | 99.9980 | Q47.08 |
| Hap2 | Chr16 | 3.368 | 1928.4 | 99.8336 | 58 | 0.0017 | 99.9983 | Q47.64 |
| Hap2 | scaffold 17 | 0.023 | 6690.1 | 99.8012 | 2 | 0.0088 | 99.9912 | Q40.54 |
| Hap2 | scaffold 18 | 0.282 | 1789.3 | 99.9993 | 11 | 0.0039 | 99.9961 | Q44.09 |

Q∞ indicates an infinite estimated quality value resulting from zero nucleotide corrections during Polypolish and does not necessarily imply an error-free assembly.

**Supplementary Table 2.** Homeodomain-like domains identified in predicted homeodomain (HD) mating-type proteins from the S1W6 Hap1 and Hap2 assemblies using the SUPERFAMILY 2.0 database.

| Assembly | Gene ID | Database | Accession | Domain | Start | End |
|---|---|---|---|---|---|---|
| Hap1 | E-HD2 | SUPERFAMILY | SSF46689 | Homeodomain-like | 192 | 237 |
| Hap1 | W-HD2 | SUPERFAMILY | SSF46689 | Homeodomain-like | 194 | 238 |
| Hap1 | W-HD1 | SUPERFAMILY | SSF46689 | Homeodomain-like | 188 | 239 |
| Hap2 | W-HD1 | SUPERFAMILY | SSF46689 | Homeodomain-like | 188 | 239 |
| Hap2 | E-HD1 | SUPERFAMILY | SSF46689 | Homeodomain-like | 183 | 234 |
| Hap2 | E-HD2 | SUPERFAMILY | SSF46689 | Homeodomain-like | 192 | 236 |
| Hap2 | W-HD2 | SUPERFAMILY | SSF46689 | Homeodomain-like | 195 | 239 |

Start and end positions indicate the amino acid coordinates of each predicted domain.

**Supplementary Table 3.** Distribution of predicted effectors and carbohydrate-active enzyme (CAZyme) classes across the two haplotypes of S1W6. Predicted effectors were classified as apoplastic, cytoplasmic, or dual. Total effectors represent the sum of all three effector classes. CAZymes were grouped into six classes: AA, auxiliary activities; CBM, carbohydrate-binding modules; CE, carbohydrate esterases; GH, glycoside hydrolases; GT, glycosyltransferases; and PL, polysaccharide lyases. Total CAZyme class counts represent the sum of assignments across all six classes, whereas total unique CAZyme proteins represent the number of distinct proteins with at least one CAZyme annotation.

| Category | Hap1 | Hap2 |
|---|---|---|
| **Predicted Effectors** | | |
| Apoplastic | 253 | 255 |
| Cytoplasmic | 87 | 84 |
| Dual | 65 | 62 |
| Total effectors | 405 | 401 |
| **CAZymes** | | |
| AA | 195 | 199 |
| CBM | 150 | 143 |
| CE | 160 | 158 |
| GH | 390 | 392 |
| GT | 240 | 221 |
| PL | 141 | 135 |
| Total CAZyme class counts | 1,276 | 1,248 |
| Total Unique CAZyme proteins | 1,152 | 1,125 |

**Supplementary Table 4.** Distribution of predicted biosynthetic gene clusters (BGCs) in the S1W6 Hap1 and Hap2 assemblies.

| Haplotype | Chromosome | BGC type | Length (bp) |
|---|---|---|---|
| Hap1 | Chr02 | terpene | 32,687 |
| Hap1 | Chr02 | terpene | 31,808 |
| Hap1 | Chr02 | terpene | 31,405 |
| Hap1 | Chr04 | terpene | 31,408 |
| Hap1 | Chr05 | terpene | 31,123 |
| Hap1 | Chr09 | terpene | 31,428 |
| Hap1 | Chr09 | terpene | 31,405 |
| Hap1 | Chr09 | terpene | 31,419 |
| Hap1 | Chr12 | terpene | 31,346 |
| Hap1 | Chr13 | terpene | 31,473 |
| Hap1 | Chr15 | NRPS | 67,905 |
| Hap1 | Chr16 | NRPS | 68,131 |
| Hap2 | Chr02 | terpene | 32,686 |
| Hap2 | Chr02 | terpene | 31,808 |
| Hap2 | Chr02 | terpene | 31,405 |
| Hap2 | Chr04 | terpene | 31,426 |
| Hap2 | Chr05 | terpene | 31,123 |
| Hap2 | Chr06 | terpene | 31,386 |
| Hap2 | Chr09 | terpene | 31,428 |
| Hap2 | Chr09 | terpene | 31,405 |
| Hap2 | Chr09 | terpene | 31,419 |
| Hap2 | Chr12 | terpene | 31,385 |
| Hap2 | Chr13 | terpene | 31,473 |

| | | | |
|---|---|---|---|
| Hap2 | Chr15 | NRPS | 67,906 |
| Hap2 | Chr16 | NRPS | 68,117 |

**Supplementary Figure 1.** Hi-C contact maps of the S1W6 Hap1 (left) and Hap2 (right) assemblies. Cyan and green outlines delineate the 16 chromosomes in Hap1 and Hap2, respectively. Color intensity represents $log_{10}$-transformed Hi-C contact frequency, with brighter colors indicating higher contact frequencies.

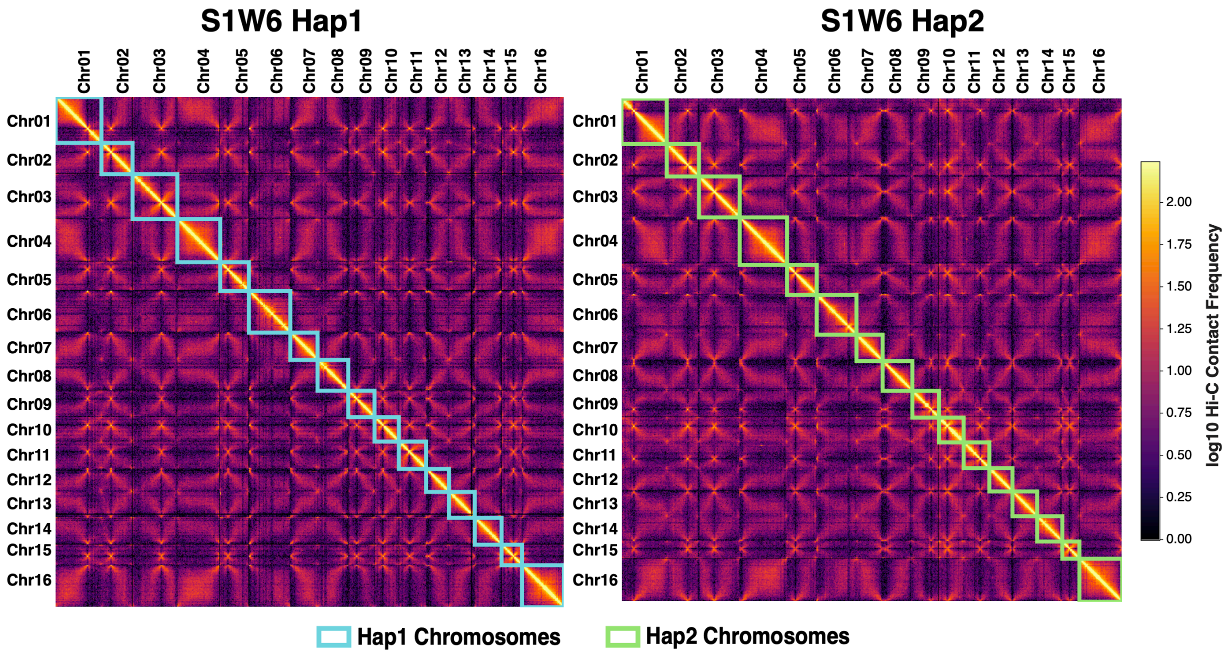

**Supplementary Figure 2.** Pairwise alignment of the Hap1 and Hap2 genome assemblies reveals collinearity between haplotypes. The asterisk represents scaffold 18 in the Hap2 assembly.

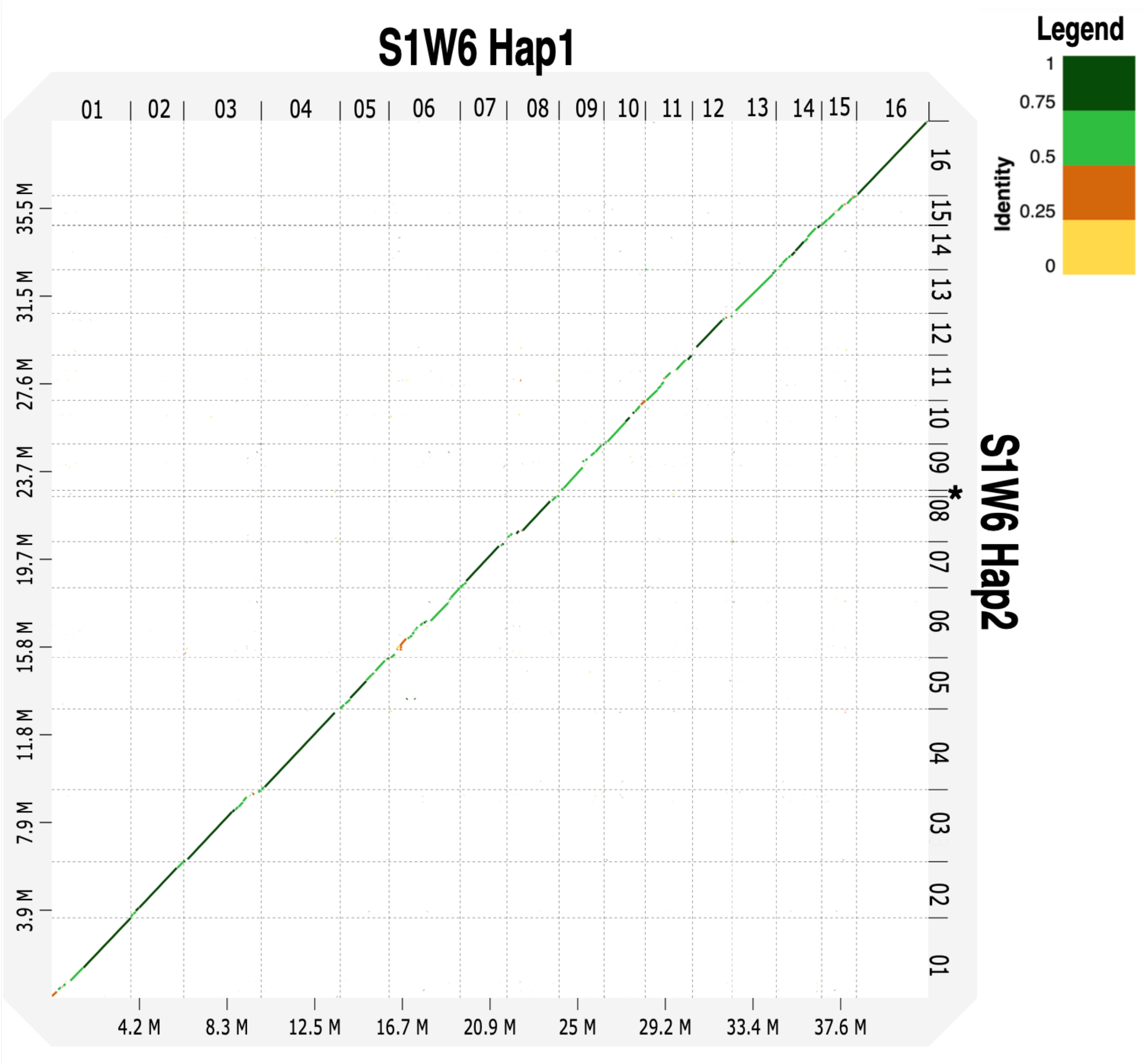

**Supplementary Figure 3.** Predicted transmembrane topology of the STE3 pheromone receptor in S1W6 Hap1 and Hap2 and AG1-IA isolate XN. Transmembrane helices were predicted using TMHMM. Posterior probabilities for transmembrane, intracellular, and extracellular regions are shown across each protein sequence.

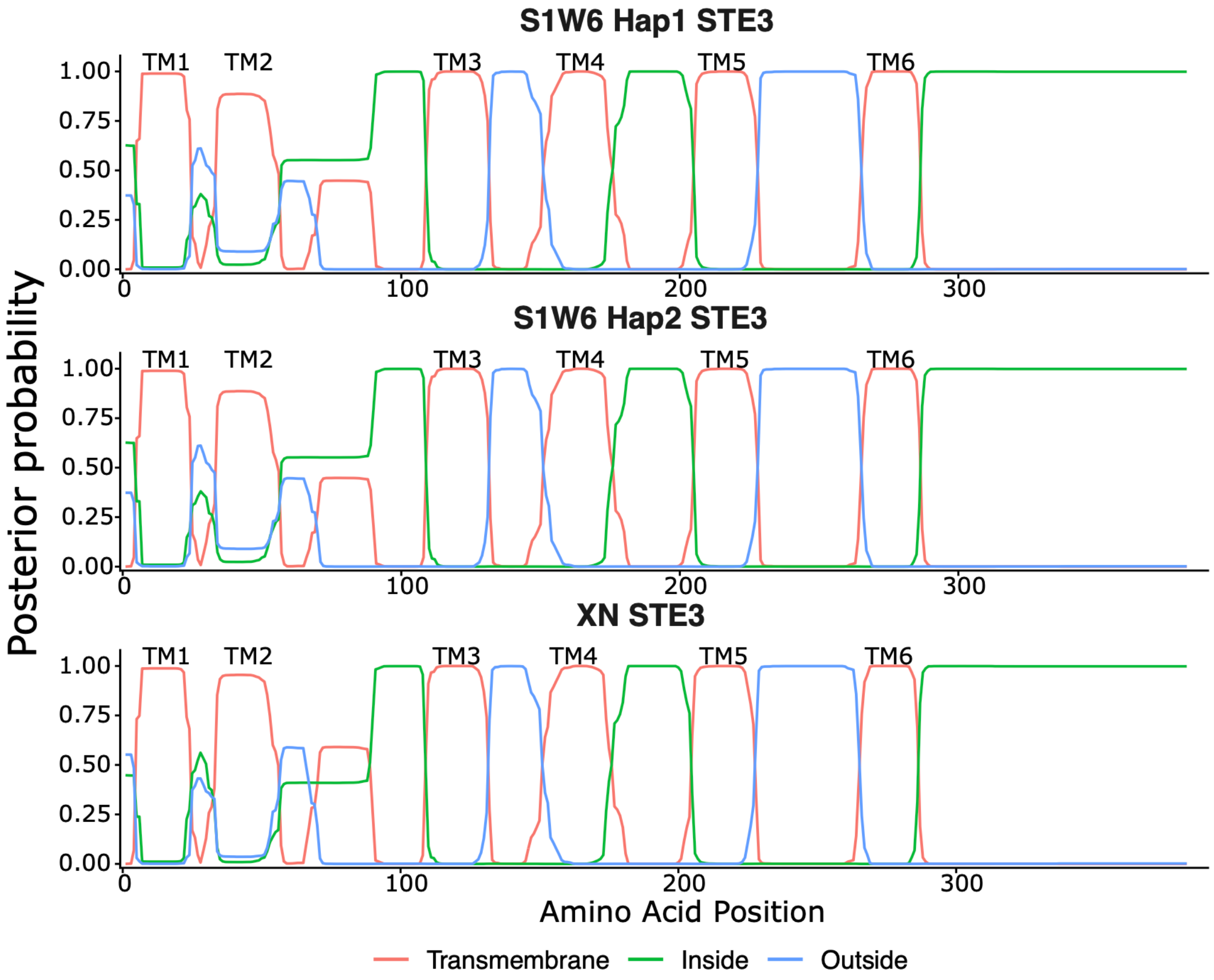
